# A predicted amphipathic helix contributes to the efficient S-acylation of the *Arabidopsis thaliana* Calcineurin B-like protein 2

**DOI:** 10.1101/2023.01.17.524453

**Authors:** Thomas Wyganowski, Linda Beckmann, Oliver Batistič

**Affiliations:** Westfälische Universität Münster; Westfälische Wilhelms Universität Münster

**Author notes:** corresponding author, Address: Institut für Biologie und Biotechnologie der Pflanzen, Universität Münster, Schlossplatz 7, 48149 Münster, Germany, Mail.

**Keywords:** amphipathic helix, palmitoylation, S-acylation, vacuolar membrane, tonoplast, CBL, protein S-acyltransferase

## Abstract

Membrane targeting of the Calcineurin B-like (CBL) calcium sensor proteins through protein S-acylation is crucial for various processes in plants, like nutrient uptake, plant development, and response to abiotic and biotic stresses. Certain CBLs target specifically to the vacuolar membrane, but which factors contribute to this particular localization and to the lipid modification efficiency are not yet known. Here, we examined the structural features of the N-terminus of *Arabidopsis thaliana* CBL2 and show that the lipid-modified cysteines are integrated within a predicted amphipathic helix. Mutations of amino acids, which contribute to the formation of this specific domain, affect S-acylation efficiency, membrane binding and function of CBL2. Interestingly, overexpression of the protein S-acyl transferase (PAT) 10 can compensate for the binding deficiency of a CBL2 mutant variant, which harbours a helix breaker mutation. This indicates that helix formation is rather involved in the S-acylation mechanism and is less important for membrane binding. Moreover, the introduction of basic residues resulted in a partial shift of the protein from the vacuolar to the plasma membrane, indicating that the underrepresentation of positively charged amino acids contributes to the vacuolar targeting specificity. Overall, our data suggest that helix formation is potentially an initial step in the S-acylation process and provides a deeper understanding of the mechanistic interplay between PATs and tonoplast targeted CBLs.

## Introduction

In all living organisms, the Calcium ion is an important second messenger regulating development, metabolic processes and the response to environmental stimuli.^1^ Spatio-temporal changes in cellular calcium levels, which can vary depending on the incoming signals, have to be converted into a protein-based signal that can be further processed by the cellular machinery. In plants, Calcium signals are decoded by the “classical” Calmodulin and related Calmodulin-like proteins (CMLs), by the family of protein kinases with calmodulin-like domains (Calcium dependent protein kinases, CDPKs), and by Calcineurin B-like (CBLs) proteins, which are related to mammalian Neuronal Calcium Sensors (NCS).^2,3^ CBLs interact with a group of Sucrose non-fermenting (Snf1) related kinases (CBL interacting protein kinases, CIPKs) and thereby regulate the activity of different membrane transporters, channels, pumps and NADPH oxidases.^4–8^ In higher plants like *Arabidopsis thaliana*, the CBLs can be functionally classified into two major groups. The first group, including CBL1, CBL4 and CBL9 localizes to the plasma membrane due to N-terminal N-myristoylation and S-acylation.^9–11^ The second group, which consists of the *Arabidopsis* CBLs 2, 3 and 6, binds to the vacuolar membrane. This targeting is mediated by a conserved N-terminal domain ^12^, which is modified by S-acylation ^13,14^ and requires the activity of *A. thaliana* protein S-acyltransferase (PAT) 10 enzyme.^15^ S-acylation, the reversible thioesterification of cysteine residues, often occurs in presence of other hydrophobic protein domains like transmembrane domains ^16^, or the vicinity of other lipid modifications like N-myristoylation ^17^, but such structures are missing in the tonoplast targeted *Arabidopsis* CBLs 2/3/6. It, therefore, remained unclear which structural features of those proteins are required to initiate S-acylation of the N-terminal domain, thereby mediating specific targeting to the vacuolar membrane.

Amphipathic helices represent distinct membrane-binding domains, where hydrophobic and hydrophilic amino acids are separated on the two opposite sides of an alpha helix. The hydrophobic side of the helix can be embedded into the lipid layer of a membrane, while the hydrophilic side is faced to the solvent. Positively charged amino acids, especially at the polar - non-polar interface of the helix, can further contribute to the membrane binding by interacting with negatively charged phospholipids and by penetrating the lipid bilayer with the hydrophobic part of the lysine sidechains (snorkelling model).^18,19^ Certain amphipathic helices can specifically bind to lipid bilayers with distinct lipid or structural profiles and thereby can target to a defined cellular membrane.^20^ Besides, the insertion of an amphipathic helix into lipid bilayers can affect the membrane curvature, thereby inducing vesicle formation and excision.^21,22^ Therefore, amphipathic helices are prominent in proteins which mediate membrane remodelling.^20,23^

Here we show that certain mutations in the N-terminal domain of *Arabidopsis* CBL2 can affect membrane binding of the protein, although the lipid modification sites were not directly altered. The introduced mutations affect the function of a predicted amphipathic helix, indicating that the formation of such a structure is important for CBL2 membrane association. Moreover, our results indicate that the effect on helix formation rather affects the S-acylation mechanism than membrane binding, and therefore can be compensated by overexpression of the respective enzyme. This work provides further insights into the structural requirements for the efficient S-acylation and specific tonoplast targeting of this certain subgroup of calcium-binding proteins and of potentially other proteins, which contain structural similar domains.

## Results

### The CBL2 N-terminus forms a predicted amphipathic helix

Previous work showed that the N-terminus of CBL2 is required and sufficient to target proteins to the vacuolar membrane and that this targeting requires S-acylation of the N-terminal cysteines C4, C12 and C18.^12,13^ However, it is still unclear why CBL2 preferentially binds to the vacuolar membrane and how this mechanism allows the disjunction from other membranes. Moreover, since CBL2 lacks other lipid modification sites or membrane-spanning domains, it is unclear how CBL2 S-acylation is mechanistically initiated. This prompted us to further analyse the structural nature of the N-terminal protein domain and to determine the requirements in membrane targeting and lipid modification of CBL2. In a previous work the crystal structure of CBL2 was resolved but the N-terminus was excluded.^24^ Therefore, we used different platforms to predict the structure of the N-terminus, like the template-based prediction programmes i-TASSER and RaptorX as well as the matrix dependent structure prediction programmes PSIPRED and PEP-FOLD3.^25–28^ Recently, during the completion of this publication, the deep learning-based protein structure prediction program alphafold also introduced a structure for CBL2 to the uniport protein database.^29–31^ When using the first 30 amino acids from CBL2, all programmes predicted a potential α-helix of the N-terminal domain, especially between the amino acids 3 and 16 (all subsequent amino acid numberings include the initiator methionine; Figure 1A). To get further insights into the potential structure, we performed a hydrophobic cluster analysis (HCA) to determine the distribution of the amino acids within the N-terminus.^32^ This revealed that hydrophobic amino acids between amino acid 5 and 19 form a potential cluster (Figure 1B). Interestingly, this cluster exhibits a rather horizontal distribution, which can indicate the presence of an amphipathic helix.^32^ Therefore, we used further prediction programmes to determine if the N-terminus of CBL2 could form an amphipathic α-helix. We used amphipaseek to determine the grade of amphipathic helix formation in CBL2 (within the first 40 amino acids), and compared this to the amphipathic helix which is formed in the M2 protein of the influenza A virus (using amino acids 43-82).^33^ This shows that M2 is forming an amphipathic helix in the first half of the sequence with relatively high probability (max. score 0,27), which corresponds to the actual amphipathic helix formed by the amino acids 45 and 62 (Supplemental Figure 1A). In contrast, amphipathic helix formation in CBL2 shows a lower degree (max. score 0,085) but is potentially formed in the first half of the analysed peptide sequence (Supplemental Figure 1A). This corresponds to the previously identified domain which is sufficient for targeting and lipid modification.^12,13^ We used Heliquest to draw a helical wheel and to determine the properties of a potential helix. Similar to the amphipathic helix in M2 (amino acids 45-62), all hydrophobic amino acids in the CBL2 N-terminus could be arranged on one side of the potential helix (amino acids 5, 8, 11, 15, 16, 19). On the other hand, all hydrophilic amino acids (amino acids 2, 3, 6, 7, 9, 10, 13, 14, 17) cluster on the other half of the helix. This clear separation of hydrophobic and hydrophilic amino acids is a hallmark of amphipathic helices (Drin and Antonny, 2010).^20^ Importantly, all cysteine residues (4, 12 and 18) in CBL2 cluster together within the hydrophobic amino acids. It was reported that cysteine residues can have a strong hydrophobic character,^34,35^ indicating that here the cysteines are part of the hydrophobic face of the amphipathic helix even in absence of the lipid modification (Figure 1; Supplemental Figure 1B). Moreover, the overall hydrophobicity of the potential helix in CBL2 is higher than compared to M2 (H= 0,688 vs. 0,519, respectively), and especially the hydrophobic moment, which can be used to estimate the degree of amphiphilicity of a helix,^36,37^ is also higher in CBL2 compared to M2 (μH= 0,547 vs. 0,474, respectively). The CBL2 N-terminus exhibits an overall neutral net charge (z= 0), while the net charge of M2 is more positive (z=+4). Moreover, the charged amino acids lysine and aspartate at positions 6 and 9 could form an “i, i+3” ion bridge, which could be involved in the stabilization of a helix.^38^

**Figure 1:**
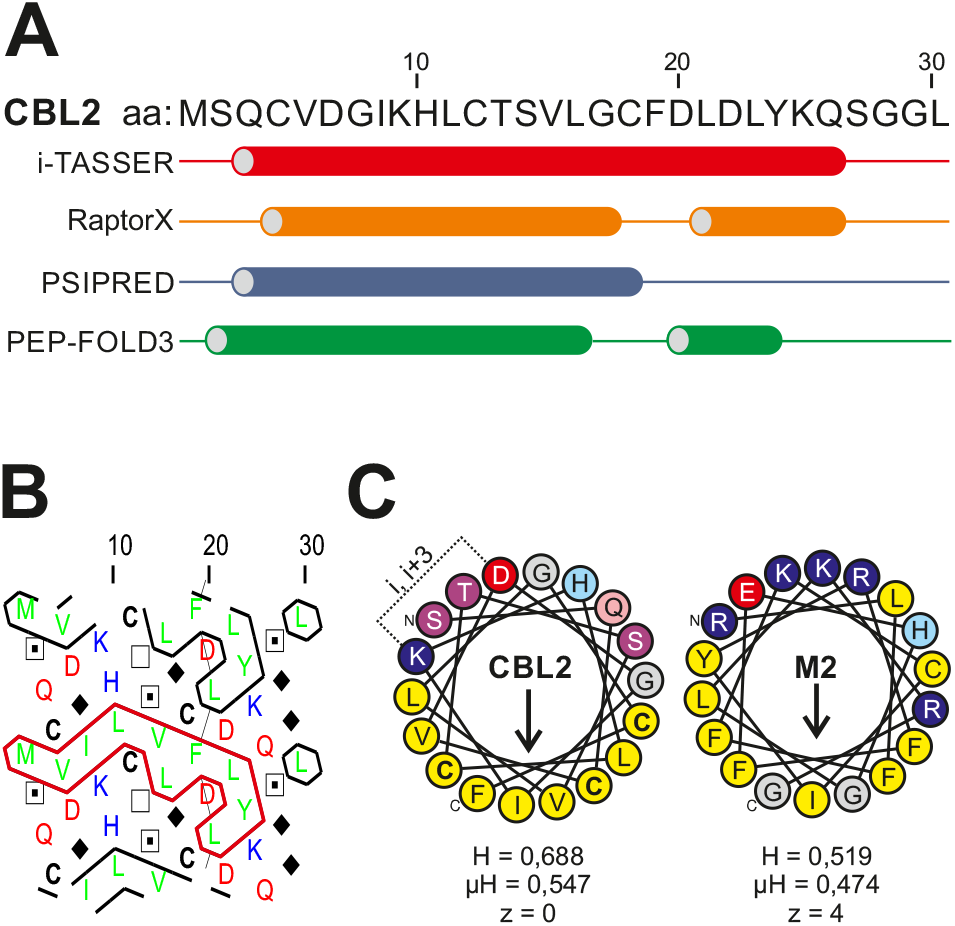
The CBL2 N-terminus forms a predicted amphipathic helix. A) Secondary structure prediction of the first 30 amino acids of *Arabidopsis thaliana* CBL2, using i-TASSER, RaptorX, PSIPRED and PEP-FOLD3. Helix formation predicted by the different algorithms is indicated by the barrels. B) Hydrophobic cluster plot of the CBL2 N-terminus (amino acids 1-30). The red bordering highlights a horizontal cluster of hydrophobic amino acids. C)Helical wheel projection of the CBL2 N-terminal amino acids 02-19 (left) and the M2 amphipathic helix (amino acids 45-62). The N-terminal and C-terminal end is indicated. Hydrophilic amino acids are clustered in the upper half, the hydrophobic amino acids (yellow colour) are clustered in the lower half of the wheel. A potential ion bridge in the CBL2 N-terminus is indicated (i, i+3). The central arrow within the wheel indicates the hydrophobic moment μH. Hydrophobicity (H), hydrophobic moment (μH) and net charge (z) values as calculated by Heliquest are given below the wheels.

Overall, these analyses revealed that the hydrophobic amino acids within CBL2 could form a functional cluster, which is typically found in amphipathic helices. Importantly, the cysteine residues arrange within this cluster, indicating that lipid modification could further contribute to the hydrophobicity and/or to the hydrophobic moment of this potential amphipathic helix.^39^ Therefore, we propose that amphipathic helix formation in the CBL2 N-terminus could be involved in membrane binding and thereby directs lipid modification of the protein. Moreover, it is likely that the specific properties of this proposed structure additionally determine the specificity in the targeting of CBL2 to the tonoplast.

### Tonoplast localization of CBL2 depends on amino acids which are potentially involved in the formation of the predicted amphipathic helix

We then wanted to address if the proposed amphipathic helix formation is involved in the targeting of CBL2 to the vacuolar membrane, besides the previously described S-acylation.^13^ For this purpose, we first examined CBL2 variants containing different mutations, which should alter the amphipathic nature of the helix but not the α-helix formation in general (Supplemental Figure 2). Mutated CBL2 variants fused to GFP were expressed in *Nicotiana benthamiana* epidermal leaf cells and the localization was compared to the co-expressed wild type CBL2 protein (CBL2^wt^), which was fused to mCherry. First, a mutation in the potential hydrophobic side was introduced to determine if this affects membrane binding. For this, valine at position 5 was replaced by the hydrophilic amino acid asparagine. In contrast to the tonoplast bound CBL2^wt^-mCherry protein, CBL2^V5N^ accumulated in the cytoplasm and nucleoplasm, indicating that hydrophobic amino acids are involved in the membrane binding of CBL2 (Figure 2A). As a control, replacing this valine by another hydrophobic amino acid like leucine did not affect the targeting of the protein to the vacuolar membrane (Figure 2B), showing that conservation of the hydrophobic character preserves membrane binding. We when introduced an asparagine mutation in the potential hydrophilic side, which should have a minor role on membrane binding. In fact, when threonine at position 13 was mutated to asparagine, this mutation did not affect the targeting of CBL2^T13N^ to the vacuolar membrane and the protein co-localized with the CBL2^wt^ protein (Figure 2C). However, we then introduced the helix breaker amino acid proline at this position (CBL2^T13P^) to determine if helix formation could play a role in CBL2 targeting. Importantly, this mutated protein accumulated in the cytosol, showing that here helix formation seems to be a prerequisite for efficient membrane binding of CBL2 (Figure 2D).

**Figure 2:**
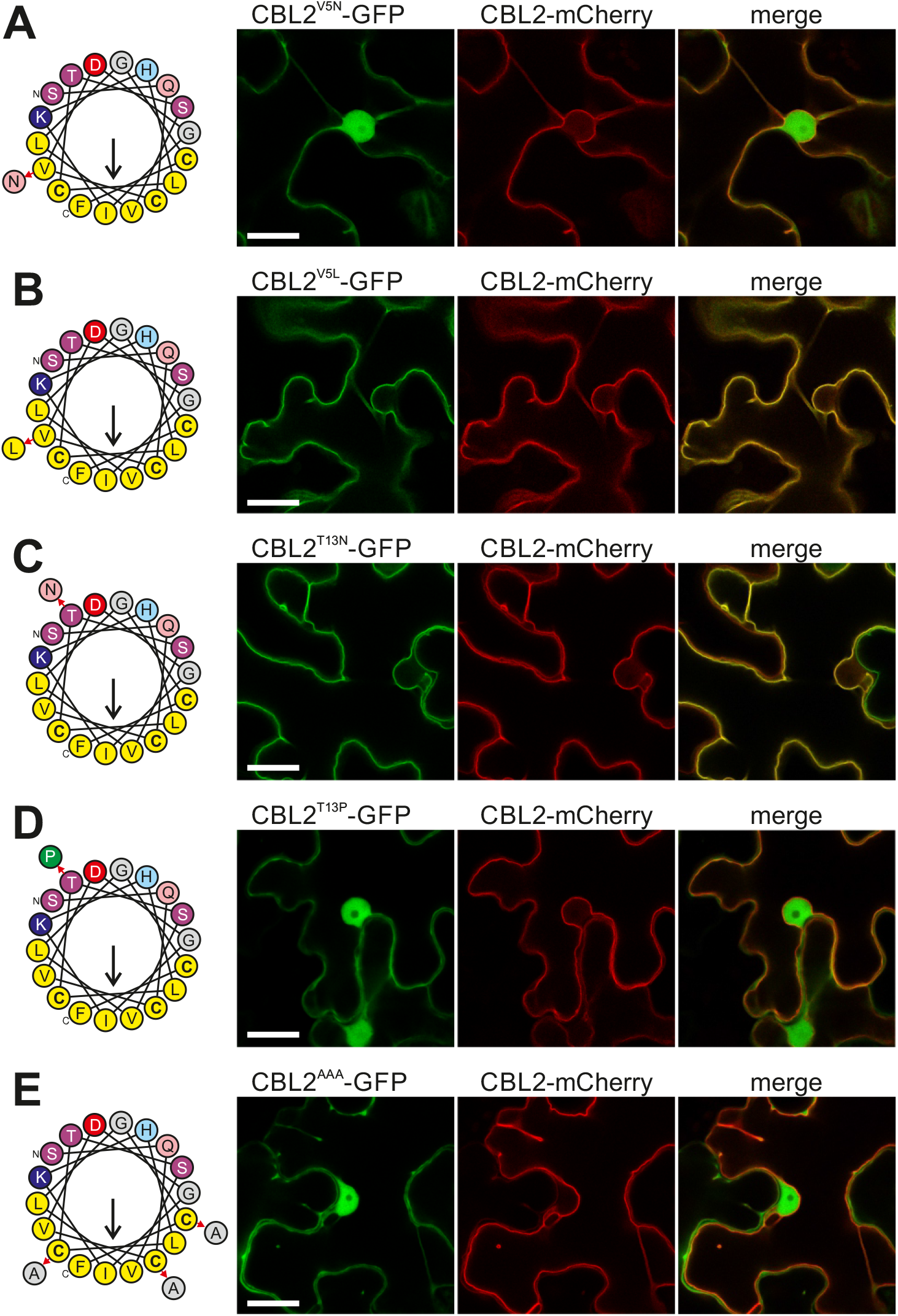
Tonoplast binding of CBL2 is affected by mutations of the potential amphipathic helix. Different CBL2 mutant versions were fused to GFP, and co-expressed with the wild type CBL2 protein fused with mCherry (CBL2^wt^-mCherry). On the left side, the helical wheel projections indicate the mutations introduced into CBL2. CBL2^V5N^ (A), CBL2^T13P^ (D) and CBL2^AAA^ (E) accumulated in the cytosol and nucleoplasm, while CBL2^V5L^ (B) and CBL2^T13N^ (C) associated with the vacuolar membrane like CBL2^wt^. The first image represents GFP, the second image represents RFP, the fluorescence is merged in the last image. Bar in the GFP image represents 20 μm.

In a previous study, it was shown that replacing cysteines with serines strongly affected membrane binding.^13^ However, this could have adversely affected the hydrophobic nature of the potential amphipathic helix. We, therefore, performed an additional experiment, where we tried to preserve a certain hydrophobicity of the amphipathic helix by replacing the cysteines 4, 12 and 18 with alanine (CBL2C4,12,18A, thereafter simplified CBL2^AAA^). However, this protein accumulated in the cytosol and did not show membrane binding in the microscopic analysis, demonstrating that intact cysteines, most likely by additional lipid modification are further required for membrane binding (Figure 2E). Finally, we performed western analysis, which revealed that all CBL2 protein variants were expressed at the expected size, showing that the differences in localizations are caused by the introduced mutations, and not by an unexpected degradation of the proteins (Supplemental Figure 2).

Overall, these microscopic localization analyses showed that other amino acids, besides the lipid-modified cysteines, are involved in the stable association of CBL2 to the vacuolar membrane. Either introducing a helix breaker in the hydrophilic side of the N-terminus or affecting the hydrophobic nature by the introduction of hydrophilic amino acids resulted in cytosolic accumulation of the CBL2 protein. This behaviour is typical for proteins which contain amphipathic helices.^40–42^ However, although this indicates the formation of an amphipathic helix, the N-terminus of CBL2 still requires lipid modifications of the cysteine residues which are positioned in the hydrophobic part of the helix and which then mediates a stable association with the tonoplast.^13^

### The CBL2^T13P^ protein functionally differs from the non-S-acylatable CBL2^AAA^ protein

To gain more insights into the requirement of the potential N-terminal helix formation we analysed the function of CBL2 variants *in planta*. CBL2, together with the highly-related CBL3 sister protein, is involved in plant development, ion homeostasis and the response to the phytohormone abscisic acid (ABA).^13,43–45^ Therefore, we determined the growth of plants, which express the three CBL2 variants (CBL2^wt^, CBL2^T13P^ and CBL2^AAA^) under the control of the endogenous *CBL2* promoter (*pCBL2* on media supplemented with CaCl_2_ (20 mM; measuring shoot fresh weight) or ABA (2 μM; measuring root length). The CBL2 variants were introduced into the *A. thaliana cbl2/cbl3* reduced-function (*cbl2/cbl3rf*) mutant, which represents a *cbl2* knock-down and *cbl3* knock-out mutant (Eckert et al., 2014). We specifically used this mutant line because we assumed that this mutant line would allow a more accurate distinction of the CBL2 variants, which would be hampered if we would have used the existing full *cbl2* and *cbl3* double knock-out plant line (named *cbl2/cbl3lf*), which is severely affected in development.^45^

First, the growth of the recombinant lines was analysed on CaCl_2_ supplemented media. Comparing WT and *cbl2/3rf* lines revealed that after 2 weeks, the shoot of the wild type plants exhibited a higher fresh weight than that of the *cbl2/3rf* mutant (Figure 3A). Interestingly, the *cbl2/3rf* lines expressing CBL2^wt^, CBL2^T13P^ and CBL2^AAA^ exhibited a gradual decline in fresh weight. *cbl2/3rf* lines with CBL2^wt^ showed highest fresh weight values, while in the *cbl2/3rf-CBL2AAA* line, we found the smallest plants with the lowest fresh weight rate. In contrast to that, the *CBL2^T13P^* lines were smaller than *CBL2^wt^* complementation lines, but were larger than the *CBL2^AAA^* plants, occupying a position between the *CBL2^wt^* and *CBL2^AAA^* plant lines.

**Figure 3:**
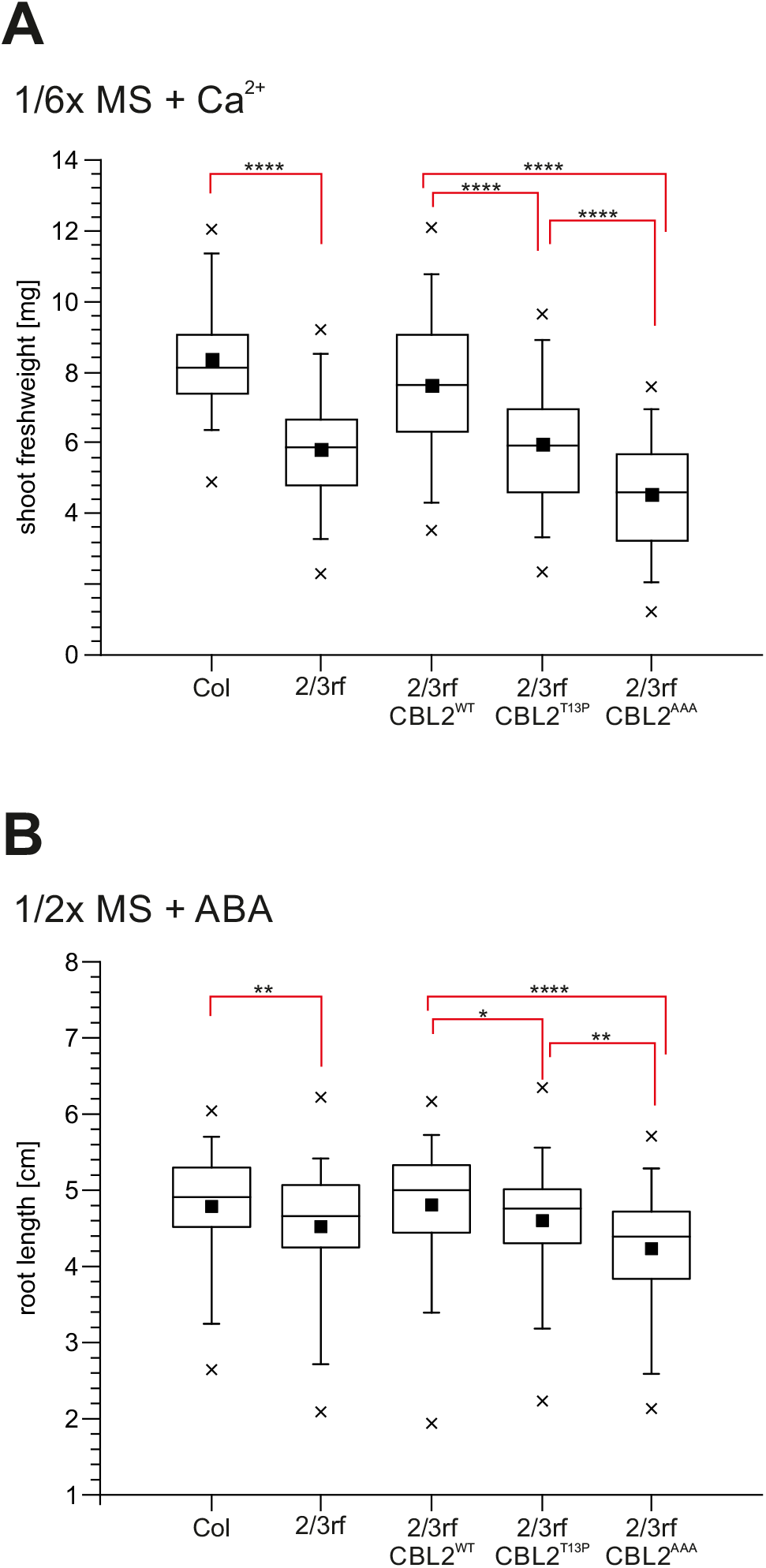
The CBL2^T13P^ mutant functionally differs from the CBL2^AAA^ variant. A) *A. thaliana* Col WT plants (WT), *cbl2/cbl3rf* (*2/3rf*), and *cbl2/cbl3rf* recombinant lines transformed with CBL2^wt^ (*2/3rf-CBL2^wt^*), CBL2^T13P^ (*2/3rf-CBL2^T13P^*) and CBL2^AAA^ (*2/3rf-CBL2AAA*) were grown on 1/6 MS media containing 20 mM CaCl2. Fresh weight of shoots was measured after 2 weeks and are depicted by box plots. Results from 3 biological samples with each at least n = 33/group were pooled and analyzed statistically by one-way ANOVA with Bonferroni’s post-hoc multiple comparisons test. B) *A. thaliana* Col WT plants (WT), *cbl2/cbl3rf* (*2/3rf*), and *cbl2/cbl3rf* recombinant lines were first germinated on 1/2x MS media for 4 days and then transferred to 1/2x MS media containing 2 μM ABA. Root extensions were measured after another 8 days of growth and are depicted by box plots. Results from 3 biological samples with each at least n = 28/group were pooled and analyzed statistically by Kruskal-Wallis with Dunn’s post-hoc multiple comparisons test. Boxes represent the 25^th^ and 75^th^ percentile and are separated by the median, while ■ specifies the mean value. Whiskers range is set at the 5^th^ and 95^th^ percentile whereas the symbol “x” represents the most distant outliers.

To get a second proof of this observation, we analyzed the root growth of these lines on ABA supplemented media (Figure 3B). Again, first, we compared the root growth (length after 12 days of germination) of wt *A. thaliana* Col-0 plants and *cbl2/cbl3rf* mutants. The *cbl2/cbl3rf* mutant plants exhibited in general shorter roots on ABA media compared to WT plants. We then determined the root length of our three *cbl2/cbl3rf-CBL2* variants. *cbl2/cbl3rf-CBL2^wt^* lines exhibited the longest roots compared to all three other lines, while the *cbl2/cbl3rf-CBL2^AAA^* lines showed the shortest roots. Again, the *cbl2/cbl3rf-CBL2^T13P^* took a middle position between the *CBL2^wt^* and *CBL2^AAA^* complementation lines.

Taken together, these experiments revealed that plants, which express *CBL2^T13P^* were functionally different compared to the non-S-acylatable *CBL2AAA* and *CBL2^wt^* lines, at least under conditions, which were tested here (shoot fresh weight on calcium-containing media, primary root growth in presence of ABA). The results indicate that a partial fraction of CBL2^T13P^ could form a functional protein and thereby enables a better growth of these plants as compared to plants expressing the non-functional CBL2^AAA^ protein version. However, the CBL2^T13P^ variant remains functionally less efficient as compared to CBL2^wt^.

### The CBL2^T13P^ mutant protein is partially bound to membranes and S-acylated

Although the microscopic analysis showed a similar cytosolic accumulation of CBL2^T13P^ and CBL2^AAA^ when both proteins are overexpressed in *N. benthamiana* leaf cells (Figure 2), the functional analysis revealed a partial better growth of CBL2^T13P^ expressing plants compared to CBL2^AAA^ (Figure 3). This points to the possibility that a certain portion of CBL2^T13P^ is able to bind to membranes, which is likely overseen in the microscopic analysis.

Therefore, to test whether CBL2^T13P^ could target to membranes, we performed subcellular fractionation analysis. For this, we replaced the GFP tag by the small HA tag, to exclude any unwanted effect of the fluorophore tag on the membrane binding efficiency of the different CBL2 protein variants. CBL2^wt^, CBL2^T13P^ and CBL2^AAA^ were transiently expressed in *N. benthamiana* cells, extracted and subsequently separated into a cytosolic/soluble and membrane-bound/insoluble fraction by ultra-high-speed centrifugation (Figure 4A). CBL2 efficiently associated with the membrane fraction while a minor fraction was present in the soluble fraction (insoluble to soluble ratio 6:1), which could represent a non-modified portion of CBL2. On the other hand, CBL2^AAA^ protein was present in the soluble fraction and not detectable in the insoluble fraction, indicating that CBL2^AAA^ is not bound to membranes. In contrast, CBL2^T13P^ was present in both the insoluble and soluble fractions and was even partially enriched in the insoluble fraction compared to the soluble sample (insoluble to soluble ratio 1,8:1). This result shows that, as already indicated by the functional analysis, CBL2^T13P^ indeed could bind to membranes, but less efficiently than the CBL2^wt^ protein.

**Figure 4:**
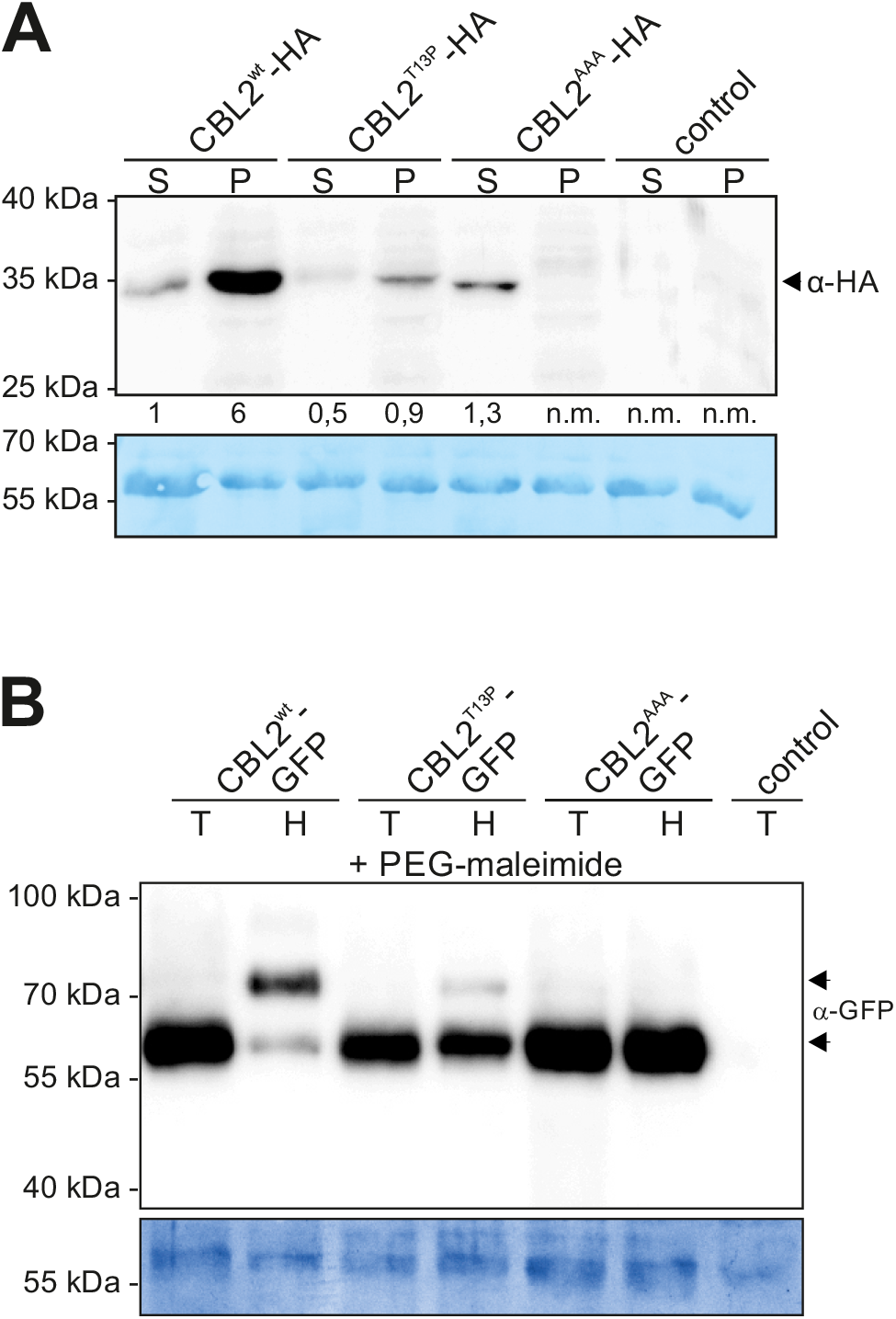
The CBL2^T13P^ helix breaker mutant is partially associated with membranes and S-acylated. A) Sub-cellular fractionation of CBL2^wt^, CBL2^T13P^ and CBL2^AAA^ (all proteins fused with HA tag). Proteins were expressed in *N. benthamiana*, and extracted proteins were separated into a soluble (S) and insoluble pellet fraction (P) by high-speed centrifugation. CBL2^wt^ and CBL2^T13P^ mainly accumulated in the pellet fraction, while CBL2^AAA^ was only detectable in the soluble fraction. The number for each lane gives the relative intensity of the bands detected (normalized to the first band [CBL2 in the soluble fraction]; n.m. = not measurable). As an antibody specificity control, CBL2^wt^-GFP was expressed and processed as described. The position of a protein marker is indicated on the left. B) S-acyl PEG switch on wild type CBL2, CBL2^T13P^ and CBL2^AAA^, all fused with GFP. Proteins were expressed in *N. benthamiana* plants, extracted in presence of NEM and subsequently incubated in either Tris control (T) or Hydroxylamine (H) solution containing PEG-maleimide. Fusion proteins were detected using a-GFP antibodies, at ~ 55 kDa. CBL2^wt^ and CBL2^T13P^ further exhibit a shifted protein in the Hydroxylamine sample at around 70 kDa, indicating S-acylation of the protein. CBL2^wt^-HA protein was expressed and used as specificity control (Tris control sample is shown). The position of a protein marker is indicated on the left. The rbcL protein was stained with Coomassie as a loading control.

However, this result also indicates that the membrane-bound fraction of CBL2^T13P^ could be still lipid-modified, but likely less efficiently than the CBL2^wt^ protein. To test this, we performed an S-acyl-PEG-switch assay, which is based on the replacement of S-acyl groups by the heavier PEG molecule and the resulting increase in protein mass detected in SDS-PAGE/Western analysis.^46,47^ We expressed CBL2^wt^, CBL2^T13P^ and the non-S-acylatable CBL2^AAA^ transiently in *N. benthamiana* cells (here using GFP fusion proteins). Protein extracts were prepared in the presence of the thiol group blocker NEM. Subsequently, thioester groups were removed by Hydroxylamine and replaced by PEG-maleimide. As shown in Figure 4B, we detected the CBL2^wt^ protein at the expected size of around 55 kDa in the Tris control (control without the hydroxylamine-mediated cleavage of the palmitoyl-thioester bond) and the Hydroxylamine sample. Importantly, in the protein extracts treated with Hydroxylamine, we detected a further shifted protein at approximately 70 kDa, indicating that S-acyl groups of CBL2 were replaced by PEG groups. As expected, the non-S-acylatable CBL2^AAA^ protein appeared as a single protein in the Tris sample as well as in the Hydroxylamine sample. In contrast to that, CBL2^T13P^ was present as a single band in the Tris sample, while a second shifted but rather weak band appeared in the Hydroxylamine sample.

These results indicate that CBL2^T13P^ can still be lipid-modified, but the introduced mutation seems to affect S-acylation efficiency and in turn by this affects membrane binding. We, therefore, speculated that increasing the efficiency of the S-acylation mechanism of CBL2^T13P^ could result in a more efficient tonoplast association. Thus, we tested the possibility to overexpress an enzyme, which is involved in CBL2 modification and thereby bypass the T13P mutation in the N-terminus. It was shown that CBL2 targeting to the vacuolar membrane is affected in the *Arabidopsis pat10* mutant ^15^ indicating that PAT10 represents the enzyme required for CBL2 S-acylation. Therefore, we analysed the localization of CBL2^T13P^ and co-overexpressed the potential CBL2 modifying enzyme *Arabidopsis* PAT10 (fused to mCherry) or a non-active PAT10^DHHA^ mutant version in *N. benthamiana* leaves (Figure 5). CBL2^T13P^ expressed alone showed the cytosolic accumulation as observed previously (Figure 5). As proposed, when PAT10 was co-expressed, we observed an efficient targeting of CBL2^T13P^ to the vacuolar membrane, which was not observed when the non-active PAT10^DHHA^ protein was co-expressed. As a further control, we also overexpressed *A. thaliana* PAT11, which is also targeted to the vacuolar membrane (Figure 5).^48^ However, overexpression of this enzyme does not mediate the targeting of CBL2^T13P^ to the vacuolar membrane, indicating that PAT10 is mediating a more efficient S-acylation even towards the mutated CBL2 substrate than compared to PAT11. These results corroborated our conclusion that CBL2^T13P^ seems to be rather affected in S-acylation efficiency, and that increasing S-acylation effectivity by overexpressing the respective enzyme can overcome the mutation effect, which finally mediates a more efficient membrane binding.

**Figure 5:**
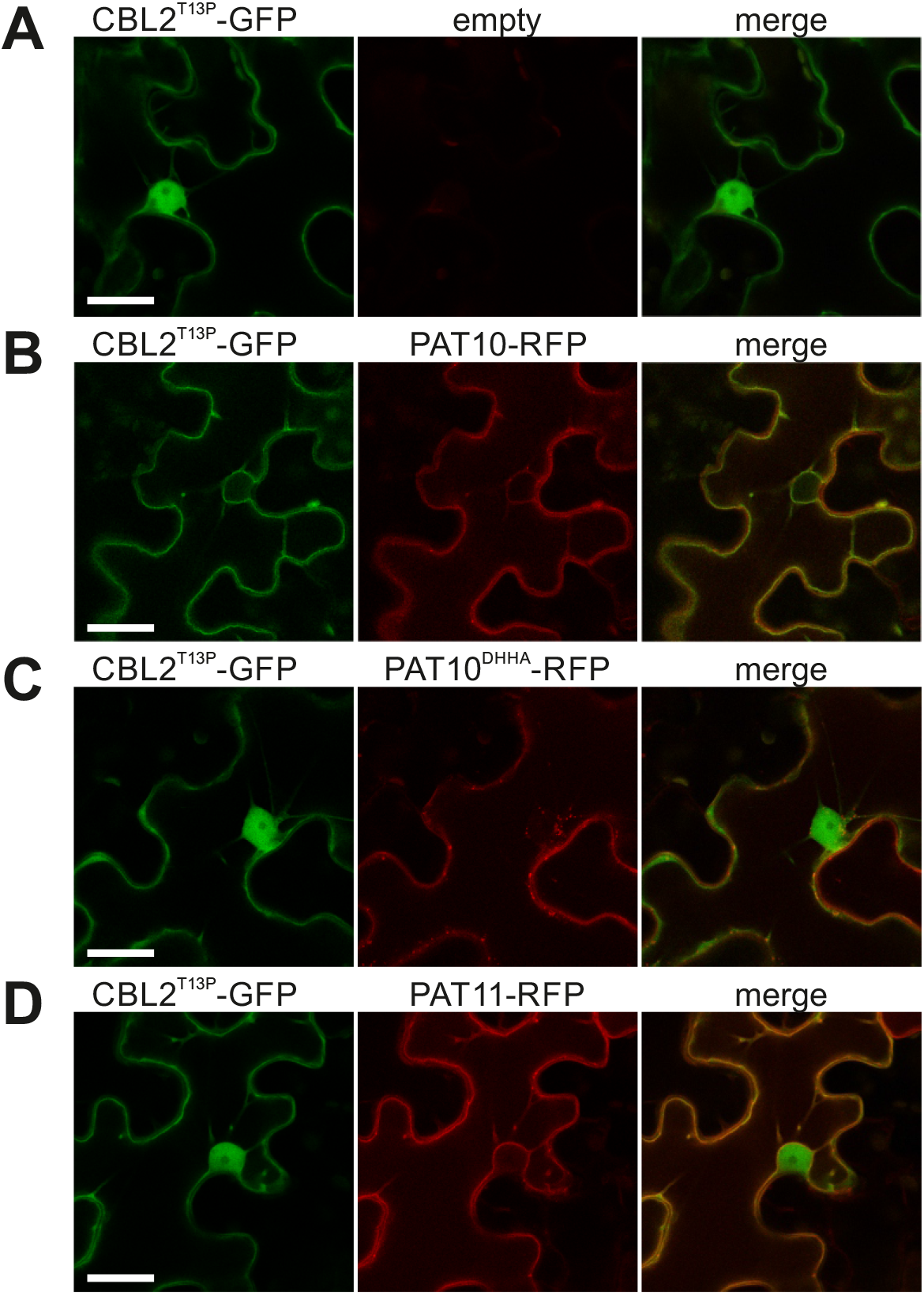
Overexpression of *Arabidopsis* PAT10 compensates the targeting deficiency of CBL2^T13P^ in *N. benthamiana*. CBL2^T13P^ was expressed in *N. benthamiana* either alone (A), together with *Arabidopsis thaliana* PAT10-mCherry (B), together with the inactive PAT10DHHA-mCherry mutant version (C) or the PAT11-mCherry protein (D). CBL2^T13P^ expressed either alone or together with PAT10DHHA or PAT11 accumulates in the cytosol, but targets to the vacuolar membrane in presence of the active PAT10 protein. The first image represents GFP, the second image represents mCherry, the fluorescence is merged in the last image. Bar in the GFP image represents 20 μm.

Overall, these analyses revealed that CBL2^T13P^ can be partially lipid-modified and that this, in turn, results in partial membrane binding of the protein. This could explain why in phenotypical analysis CBL2^T13P^ is still functional, at least partially compared to the non-functional CBL2^AAA^.

The fraction, which is bound to membranes, could be sufficient to mediate the required threshold for CBL2 function. Moreover, these results further indicate that after lipid modification of the N-terminus helix formation seems to become less important for membrane binding, at least when the helix breaking mutation is introduced at the position used here.

### The lack of basic residues in the CBL2 N-terminus contributes to the targeting specificity to the vacuolar membrane

All in all, these results implied that helix formation in the CBL2 N-terminus is rather required for efficient S-acylation, but it remained unclear if and how the N-terminus also affects targeting specificity. In fact, it cannot be fully excluded that amphipathic helix formation is required for an initial membrane binding to the desired membrane and into the vicinity of the respective enzyme before CBL2 gets lipid-modified. Normally, basic residues can contribute to the binding efficiency of amphipathic helices, but we noted that the CBL2 N-terminus is rather poor in basic amino acids (Figure 1C). We, therefore, questioned if the lack of basic residues in CBL2 somehow contributes to the targeting of the protein. We replaced several of the amino acids in the hydrophilic side by basic amino acids, especially the ones which are spatially at the amphiphilic interface of the helix (serine 2, glycine 7 and serine 14, changed to lysine). This protein was co-expressed with CBL2-mCherry in *N. benthamiana*. Importantly, CBL2^S2KG7KS14K^ did not accumulate in the cytosol, showing that membrane binding was rather unaffected. However, although CBL2^S2KG7KS14K^ was targeted to the vacuolar membrane like the CBL2^wt^ protein, a certain fluorescence fraction was observed not co-localizing with CBL2^wt^ (Figure 6A). Further investigations using a plasma membrane marker protein (TM23-RFP) ^49^ revealed that CBL2^S2KG7KS14K^ was partially directed to the plasma membrane (Figure 6B). To rule out that the mutations *per se* and not the specific introduction of the basic residues mediated the partial mistargeting of CBL2^S2KG7KS14K^, we additionally tested the localization of a CBL2 variant in which the respective amino acids were replaced by alanine residues (CBL2^S2AG7AS14A^). However, this protein variant was not targeted to the plasma membrane and efficiently localized to the tonoplast (Figure 6C).

**Figure 6:**
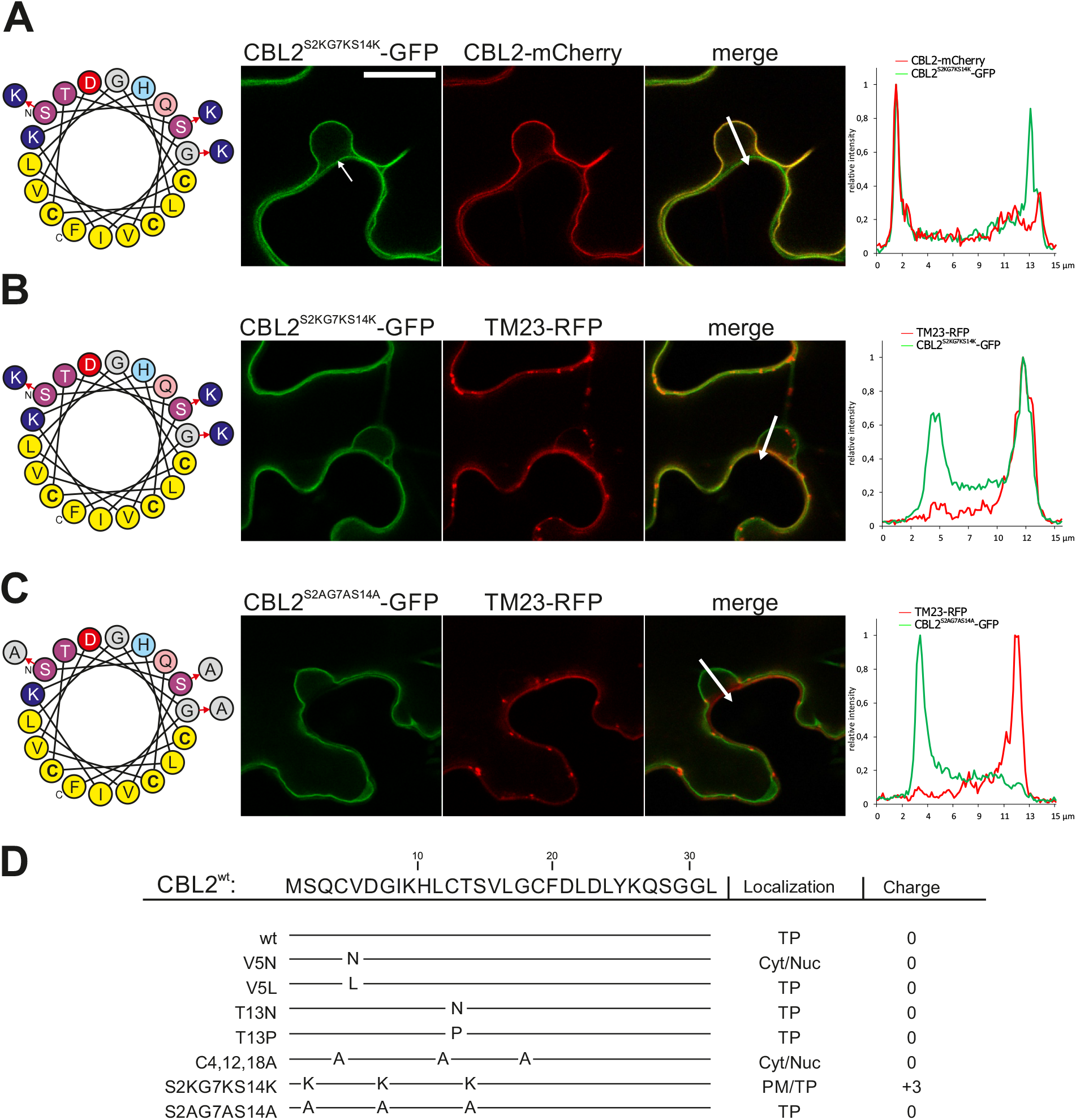
Introducing basic residues in CBL2 results in partial plasma membrane targeting. A) Serine2, glycine7 and serine14 in CBL2 were replaced by basic amino acids (indicated in the helical wheel projection on the left). This protein CBL2^S2KG7KS14K^-GFP localizes to the vacuolar membrane as CBL2^wt^-mCherry protein but partially targets to the plasma membrane as well (indicated by the arrow in the GFP image). The fluorescence intensity was measured within the region of interest indicated by the arrow in the merged image (size of arrow/ROI represents 15 μm). The intensity profile is shown on the right. B) This additional fluorescence originating from CBL2^S2KG7KS14K^-GFP co-localizes with a plasma membrane marker TM23-RFP. The fluorescence intensity was measured within the region of interest indicated by the arrow in the merged image (size of arrow/ROI represents 15 μm). The intensity profile is shown on the right. C) As a control serine2, glycine7 and serine14 in CBL2 were replaced by alanine residues (indicated in the helical wheel projection on the left). The CBL2^S2AG7AS14A^-GFP fusion is efficiently targeted to the vacuolar membrane and does not co-localizes with the plasma membrane marker TM23-RFP. The fluorescence intensity was measured within the region of interest indicated by the arrow in the merged image (size of arrow/ROI represents 15 μm). The intensity profile is shown on the right. D) Summary of the CBL2 mutants generated in this study and their corresponding subcellular localizations. PM, Plasma Membrane; TP, Tonoplast; Cyt, Cytosol; Nuc, Nucleoplasm. The total electrostatic charge of each N-terminus is indicated.

This shows that the introduction of positively charged residues results in a partial targeting of the protein to the plasma membrane, indicating that the neutral net charge of the CBL2 N-terminus in fact somehow contributes to the targeting specificity to the tonoplast. However, the introduction of basic residues in the CBL2 N-terminus was not sufficient to completely direct the protein to the plasma membrane, indicating that other factors still promote the targeting to the tonoplast.

## Discussion

Plant cells are characterized by the presence of a large central vacuole, which can occupy up to 90% of the total cell volume.^50,51^ Besides establishing a large hydrostatic pressure (turgor) which provides stability and shape, the central vacuole is further involved in storage, detoxification and signalling processes. In particular, due to the large storing capacity for the calcium ion, the vacuole represents a central organelle within the calcium signalling machinery of land plants.^3,52^

During the evolution of plants, especially through the transition to multi-cellularity and land colonization, a profound transformation of the calcium signaling machinery system has occurred which also involved the CBL calcium sensor protein family.^2,53^ Original CBLs probably possessed a “classical” N-terminal myristoylation site, which is already present in the prototypical CBLs of non-plant protist organisms and which is also common to the related NCS proteins.^54,55^ An additional S-acylation site was then introduced in early unicellular algae, likely to enable stable plasma membrane targeting.^10,53^ However, proteins which can be phylogenetically classified as “tonoplast type” CBLs, like CBL2 or CBL6, are absent in algae and appear at earliest in Bryophytes, indicating that the evolution of this specific type of calcium sensors potentially occurred during the land colonization of plants.^2,53^ During this transition, also the central vacuole has probably become more and more important as a signalling organelle, like for example in long-distance signalling events of the multicellular organisms.^56,57^ This also required to establish signalling processors which directly act at the vacuolar membrane like the “tonoplast type” CBLs.^12,53^ For this, the proteins were equipped with an N-terminal lipid modification site, which allowed to move the CBL-CIPK signalling complex from the plasma membrane to the vacuolar membrane and are thus involved in various plant processes like development and ion homeostasis.^13,43–45,58^

Up to now, it is not fully clear how this N-terminal domain is recognized by protein S-acyl transferases and how this domain contributes to the specific targeting of the tonoplast-type CBL proteins. Protein S-acylation does not require a specific signal sequence as N-myristoylation or C-prenylation, but certain structural features contribute to the recognition by PATs, like other lipid modifications or transmembrane domains.^59^ However, such features are lacking in CBL2. Therefore, we started to analyse the structural requirements of the CBL2 N-terminus, which could contribute to the efficient lipid modification and consequently for stable association especially with the vacuolar membrane. In particular, the predicted clustering of hydrophobic amino acids, the hydropathy, and the hydrophobic moment of the CBL2 N-terminal region are features, which are typically found in amphipathic helices. In accordance to that, introducing mutations (summarized in Figure 6D), which either affected the hydrophobic side (CBL2^V5N^) or disrupted the formation of the helix (CBL2^T13P^), both influenced the membrane binding of CBL2 when overexpressed in plant cells, indicating that an amphipathic helix is involved in the targeting of CBL2. At the moment, we cannot fully explain the exact mechanism. On the one hand, the N-terminal domain could allow a direct, initial membrane binding, which is then stabilized by lipid modifications. On the other hand, the N-terminus could be involved in protein-protein interaction mechanisms to the respective lipid-modifying enzyme and therefore plays a rather indirect role in membrane binding.^60^ Moreover, the formation of a specific N-terminal structure could be also relevant to create a favourable environment to increase the reactivity of the cysteine residues and thereby promotes the S-acylation of CBL2.^61^ Which of the potential factors, or even if all three factors play a role, remains open at this time. Nevertheless, the introduction of a helix breaker mutation like in CBL2^T13P^ results in a less efficient lipid modification and reduced membrane binding, which is reflected in a reduced *in planta* function. However, the lipid modification and targeting deficiency can be compensated by the overexpression of the active *Arabidopsis* PAT10 enzyme, rather indicating that the N-terminus plays a minor role in direct membrane binding *per se*, at least after successful lipid modification and stable anchoring of the protein to the tonoplast membrane.

A further obvious feature of the CBL2 N-terminus is the lack of basic residues, which are normally often found in amphipathic helices like in M2, GRK5, or human RGS2.^33,62,63^ The basic residues can contribute to the membrane-binding stability and the targeting specificity by interacting with acidic lipid headgroups.^19,62–64^ When basic residues were introduced into the CBL2 N-terminus, this resulted in partial re-targeting of the protein to the plasma membrane. On the one hand, the basic residues could mediate the specific binding of the modified N-terminus of CBL2 to certain phospholipid species which are enriched in the plasma membrane, like that of phosphatidylinositol 4-phosphate and phosphatidylinositol (4,5)-bisphosphate.^65^ Moreover, we cannot fully exclude that other lipids are contributing to the binding of CBLs, either of the native CBL2 to the tonoplast or of the modified CBL2^S2KG7KS14K^ to the plasma membrane. As PATs are composed of transmembrane domains, the enzymes could recruit specific, energetically favorable lipids in the surrounding annular shelf, which is potentially recognized by the N-terminal CBL2 domain.^66,67^ Thereby, the specificity of the lipid modification mechanism would be determined by the combination of the enzyme, its surrounding lipids and by the target. However, we should not oversee the possibility that the introduced basic amino acids maybe enabled direct protein-protein interactions with plasma membrane-bound proteins, and thereby resulted in the partial targeting of the CBL2^S2KG7KS14K^ protein to the plasma membrane as observed. Alternatively, the basic residues of CBL2^S2KG7KS14K^ could increase the substrate specificity of PM-localized PATs towards this CBL2 mutant protein. Overall, these results showed that the lack of positive charges somehow contributes to the targeting specificity of *Arabidopsis* CBL2, and likely of the related CBL proteins 2 and 6 to the vacuolar membrane.

Finally, the properties of the CBL2 N-terminus are reminiscent for amphipathic helices present in Sar1 and Arf1 G-proteins, two prominent proteins which are involved in vesicle formation. The membrane-binding domains of these G proteins are at the very N-terminus and neutral in charge.^20^ Importantly, these Sar1/Arf1 amphipathic helices have to bind first to a flat membrane before the proteins induce vesicle formation by the recruitment of additional accessory proteins.^20,68^ In this context, it should be noted that Arf1 also employed a lipid modification (N-myristoylation) for stabilizing membrane anchoring.^69^ Interestingly, it was suggested that the lipid modification of Arf1 could be important for the integrity of the Golgi apparatus.^70^ The integration of amphipathic helices into a lipid bilayer can affect the packaging of the surrounding lipids and can result in a strong bending of the membrane.^71^ It was assumed that the integration of the rather short amphipathic helix together with the myristoyl group has less effects on the Golgi membranes than the integration of a large amphipathic helix with bulky, hydrophobic amino acids, which would otherwise result in disturbance of the surrounding lipids and the integrity of the Golgi membrane.^68^ Moreover, also the lack of basic residues could further contribute to the integrity of the membrane, as basic residues rather promote the curvature of a membrane.^72^ In this regard, the plant tonoplast rather represents a flat membrane due to the high concentration of Phosphatidylcholine ^73^, which favours the formation of flat monolayers.^74^ This opens the question if therefore CBLs employed S-acyl groups or S-acylated amphipathic helices for membrane anchoring instead of a membrane curvature inducing amphipathic helix, even though the prerequisites for such a membrane-binding domain already existed. The integration of the CBL2 N-terminus into the lipid bilayer by S-acylation on one hand could promote binding to a flat membrane and besides this minimizes the impact on the vacuolar membrane integrity which otherwise could be affected if proteins with large amphipathic and basic helices are bound to the tonoplast. Interestingly, in accordance with this assumption was a recent finding on the tonoplast targeting of the homotypic fusion and vacuolar protein sorting (HOPS) subunit VPS41 from *Arabidopsis*. This protein binds to the vacuolar membrane in a Wortmannin sensitive manner, indicating that tonoplast binding is mediated by interaction with phosphoinositides. It was noted that in contrast to the yeast VPS41 protein, Arabidopsis VPS41 lacks the amphipathic lipid packing sensor (ALPS) motif, which specifically binds to curved membranes.^75^ Of course, the lack of the ALPS motif in *Arabidopsis* VPS41 on one hand could explain the ability of VPS41 to bind to flat membranes. But in addition to that, the lack of the ALPS motif could be further relevant to minimize the disturbing effect on the lipid bilayer of the large and flat central vacuole after the membrane fusion event. Taken together, this work showed that certain amino acids in the N-terminus of CBL2, other than the lipid-modified cysteines, contribute to the efficient S-acylation and thereby to the membrane binding and tonoplast targeting of this protein sub-group. These amino acids are likely involved in the formation of a potential amphipathic helix, which seems to be a prerequisite for the lipid modification mechanism. This structure could either initiate a direct membrane binding or alternatively promote a protein-protein interaction with the S-acylating enzyme. Recent works on the structure determination of S-acyl transferases together with palmitoyl-CoA^76,77^ might also help to address the enzyme-substrate binding mechanism between CBL2 and PAT10. Hence, in the future, it will be interesting to understand the exact interplay between helix formation and protein S-acylation of this specific motif regarding the true targeting mode to the vacuolar membrane.

## Materials and Methods

### Cultivation and growth analysis of *Arabidopsis thaliana* plants

*Arabidopsis thaliana* cv Columbia-0 plants and *A. thaliana cbl2/cbl3rf* mutants ^45^ were used in this work and, except for the stress treatments indicated, cultivated in a growth chamber under a 16-h-light/8-h-dark cycle with 70% atmospheric humidity and a 22°C (day)/18°C (night) temperature regime. To generate *A. thaliana cbl2/cbl3rf:CBL2* variants, the open reading frames from *CBL2^wt^, CBL2^T13P^* and *CBL2AAA* were placed behind the *CBL2* promoter within the pGPTVII plasmid backbone.^13,78^ The plasmid was transformed into *Agrobacterium tumefaciens*, and *cbl2/cbl3rf* mutant plants were transformed by the floral dip method.^79^ The transformed lines were propagated to T2 to receive homozygous lines. The recombinant status of the plant lines was verified by PCR and sequencing of genomic DNA and the transcription of the constructs was verified by RT-PCR. The sequence of all primers used is given in the Supplemental Table 1.

For growth tests on Calcium containing media, 1/6xMS media was prepared and supplemented with 20 mM CaCl2.^44^ The seeds were surface sterilized and directly sown on the media, stratified for two days, and the plants were grown for 2 weeks in a growth chamber. For ABA root growth analysis, seeds were sterilized and placed on Agar plates containing 1/2xMS, stratified for 3 days, and germinated for 4 days under a 12-h-light/12-h-dark-18°C/22°C light/temperature cycle in a growth chamber. After germination seedlings were transferred on 1/2xMS supplemented with 2 μM ABA and post transfer root elongation was measured after another 8 days. All phenotypic analyses were performed using at least two independent lines for each variant, and analyses were performed at least three times (with freshly prepared media and freshly sterilized seeds). To determine differences of the obtained mean values, one-way ANOVA followed by post hoc Bonferroni’s multiple comparisons test for normal distributed data and Kruskal-Wallis followed by post hoc Dunn’s multiple comparisons test for data that do not follow normal distribution was performed. Statistical significance is indicated by: *P ≤ 0.05, ***P ≤ 0.001, ****P ≤ 0.0001.

### Bioinformatic analysis

The secondary protein structure was analysed using public, web-based programmes: i-TASSER ^25^, RaptorX ^26^, PSIPRED ^27^, PEP-FOLD3 ^28^, Heliquest ^80^, HCA plot analysis ^81^ and amphipaseek.^82^

### Microscopic analysis

For microscopic analysis, proteins were transiently expressed via *A. tumefaciens* mediated transient transformation of *N. benthamiana* leaves. *N. benthamiana* plants were cultivated in a greenhouse under a 12-h-light/12-h-dark cycle with 60% atmospheric humidity at 20°C. An overnight culture of Agrobacteria was resuspended in activation media (10 mM MES/KOH pH 5,8, 10 mM MgCl_2_, 150 μM Acetosyringone) and incubated for 3h at room temperature. Agrobacteria were infiltrated in fully developed leaves with a needleless syringe and infiltrated leaves were microscopically inspected 2 days after infiltration. Pictures were acquired using an inverted Leica DMI6000 microscope equipped with the Leica TCS SP5 confocal laser scanning device (Leica Microsystems). Detection of fluorescence was performed as follows: GFP, excitation at 488 nm (Ar/Kr laser) and scanning at 500 to 535 nm; RFP/mCherry, excitation at 561 nm and scanning at 565 to 595 nm. All images were acquired using a Leica 63×/1.20 water immersion objective (HCX PL Apo CS). Images were further processed using the Leica AF Lite software and Corel Photo-Paint.

### S-Acyl PEG switch and sub-cellular protein fractionation

To analyze the lipid modification of GFP tagged CBL2 protein versions, the proteins were expressed transiently in *N. benthamiana* as described above. Proteins were extracted using ~200 mg of leaf material, which was frozen in liquid nitrogen and ground (using a Mixer Mill). Proteins were extracted in 1x phosphate-buffered saline (PBS) pH 7.4 ^83^, containing 5 μl/ml protease inhibitor cocktail (PIC) (Sigma Aldrich GmbH, München, Germany), 1 mM EDTA, 1% Triton X-100, 1 % sodium deoxycholate, 0,1 % sodium-dodecyl-sulfate (SDS) and 25 mM N-ethylmaleimide (NEM), and incubated for 25 min at 50 °C and 1500 rpm in a shaker. The protein extract was cleared from the cell debris by centrifugation (2x 5 min, 2000 g, 4°C) and proteins were precipitated by methanol/chloroform to remove NEM.^84^ The protein pellet was resuspended in 12 μl/mg protein of resuspension buffer (1x PBS pH 7.4 containing 8 mM Urea and 5 % SDS). An equal amount of resuspended proteins were either incubated with Hydroxylamine (800 mM, pH 7.4) or with Tris (45 mM, pH 7.4), containing 1 mM EDTA, 2 μl/ml PIC and 1 mM methoxy-polyethyleneglycol(PEG)-maleimide (5 kDa), for 40 min under continuous rotation and subsequently precipitated using Methanol/Chloroform. The proteins were resuspended in SDS loading buffer, the protein concentration was determined and equal protein amounts were separated by SDS-PAGE and detected after western blotting. GFP tagged proteins were detected using an anti-GFP antibody (1:4000; mAb rat 3H9; Chromotek) and a secondary anti-rat antibody (1:15000; pAb goat; Life Technologies) conjugated with horseradish peroxidase. Detection was performed by an enhanced chemiluminescence reaction. Sub-cellular protein fractionation was performed as described.^10^ Briefly, proteins were expressed in *N. benthamiana*. Proteins were extracted and separated into a soluble and insoluble/membrane fraction by ultra-centrifugation (100000g, 1h, 4°C). The separation of the proteins was subsequently analyzed by western blotting. HA-tagged proteins were detected using an anti-HA antibody (1:4000; mAb mouse HA.11 MMS-101P; Covance) and a secondary anti-mouse antibody (1:15000; pAb goat; Bio-Rad) conjugated with horseradish peroxidase. Detection was performed by an enhanced chemiluminescence reaction.

## Acknowledgements

This work was supported by the Deutsche Forschungsgemeinschaft (DFG BA4742/1-1 and 1-2).

## Supplemental Figures

**Supplemental Figure 1.**
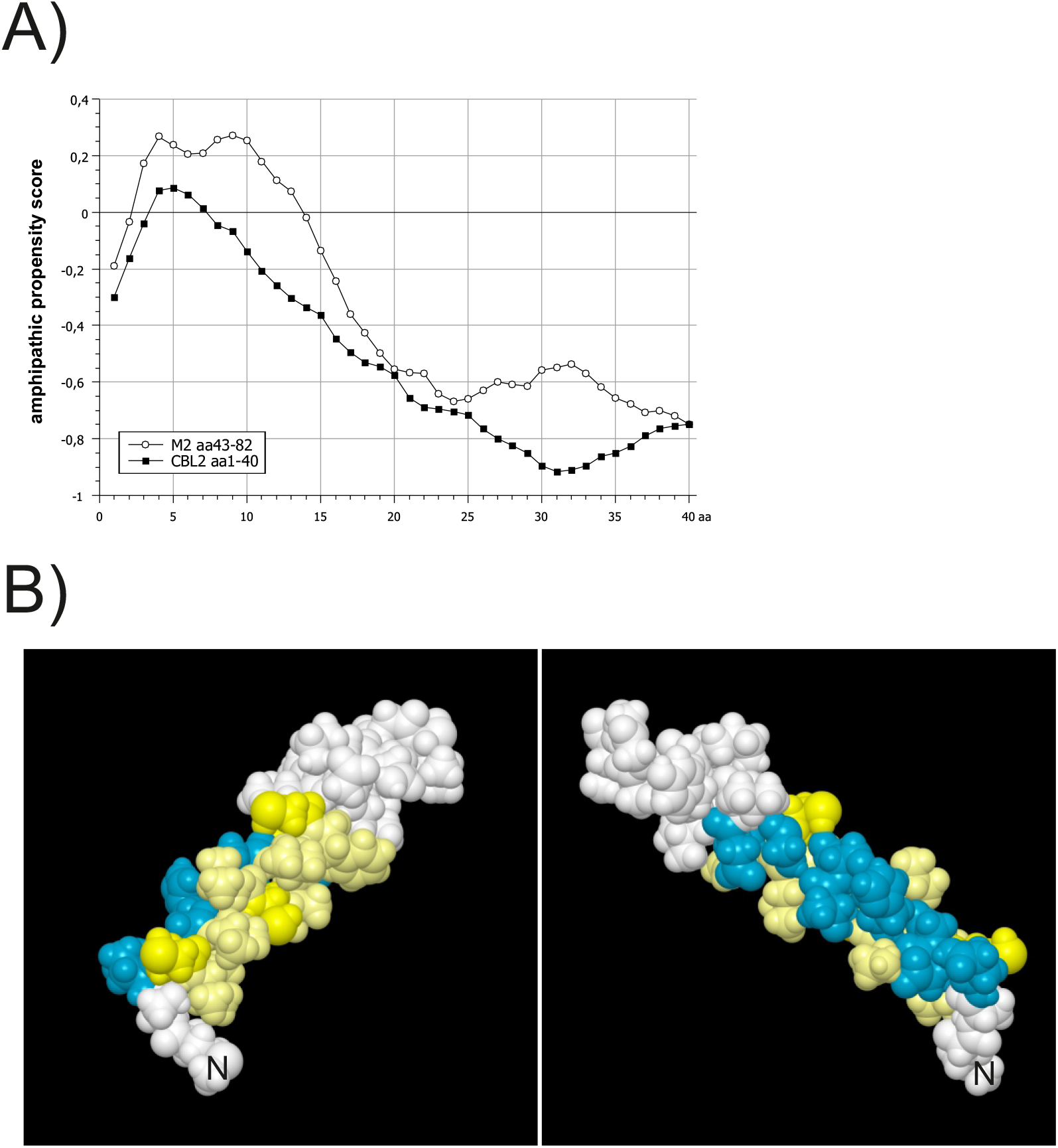
A) Amphipathic helix score of the M amphipathic helix (amino acids 43-82) and of the CBL2 N-terminus (amino acids 1-40), determined by amphipaseek. B) Space-filling model of the CBL2 N-terminus from two different angles. Amino acids 2-19 are coloured. Hydrophobic residues and Cysteines are shown in light yellow and yellow, respectively. Hydrophilic amino acids are depicted in blue. The N-terminal initiator Methionine is indicated (N).

**Supplemental Figure 2.**
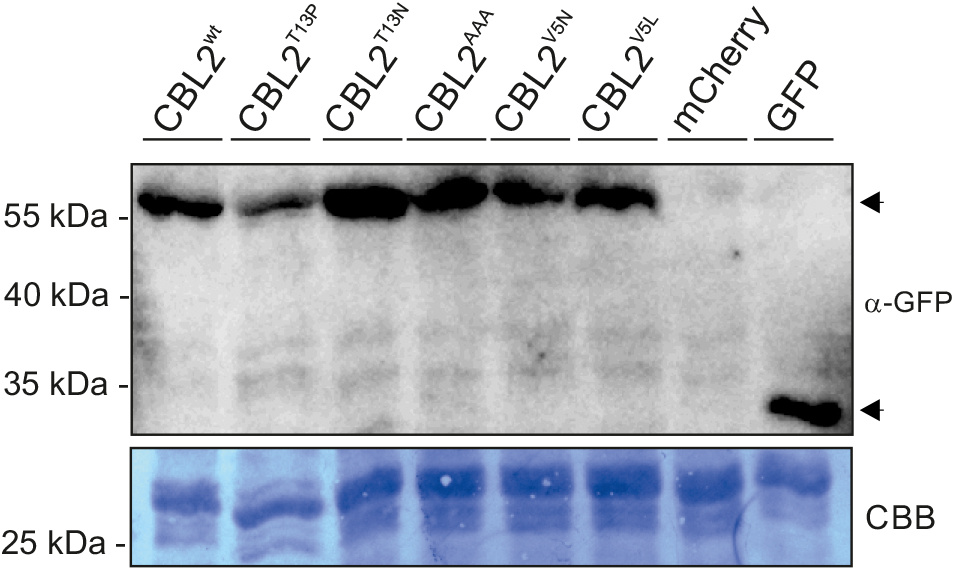
CBL2 variants are expressed as a single protein fusion. To determine the integrity of all used CBL2 variants, CBL2^wt^, CBL2^V5N^, CBL2^V5L^, CBL2^T13N^, CBL2^T13P^, and CBL2^AAA^ (all GFP fusions) were expressed in *N. benthamiana*, extracted, and analysed by western blot analysis. As an antibody specificity control, mCherry was expressed (lane 7). As a further control, unmodified GFP was expressed (last lane). The size of the proteins is indicated on the left. A loading control stained with Coomassie is shown below.

**Supplemental Figure 3.**
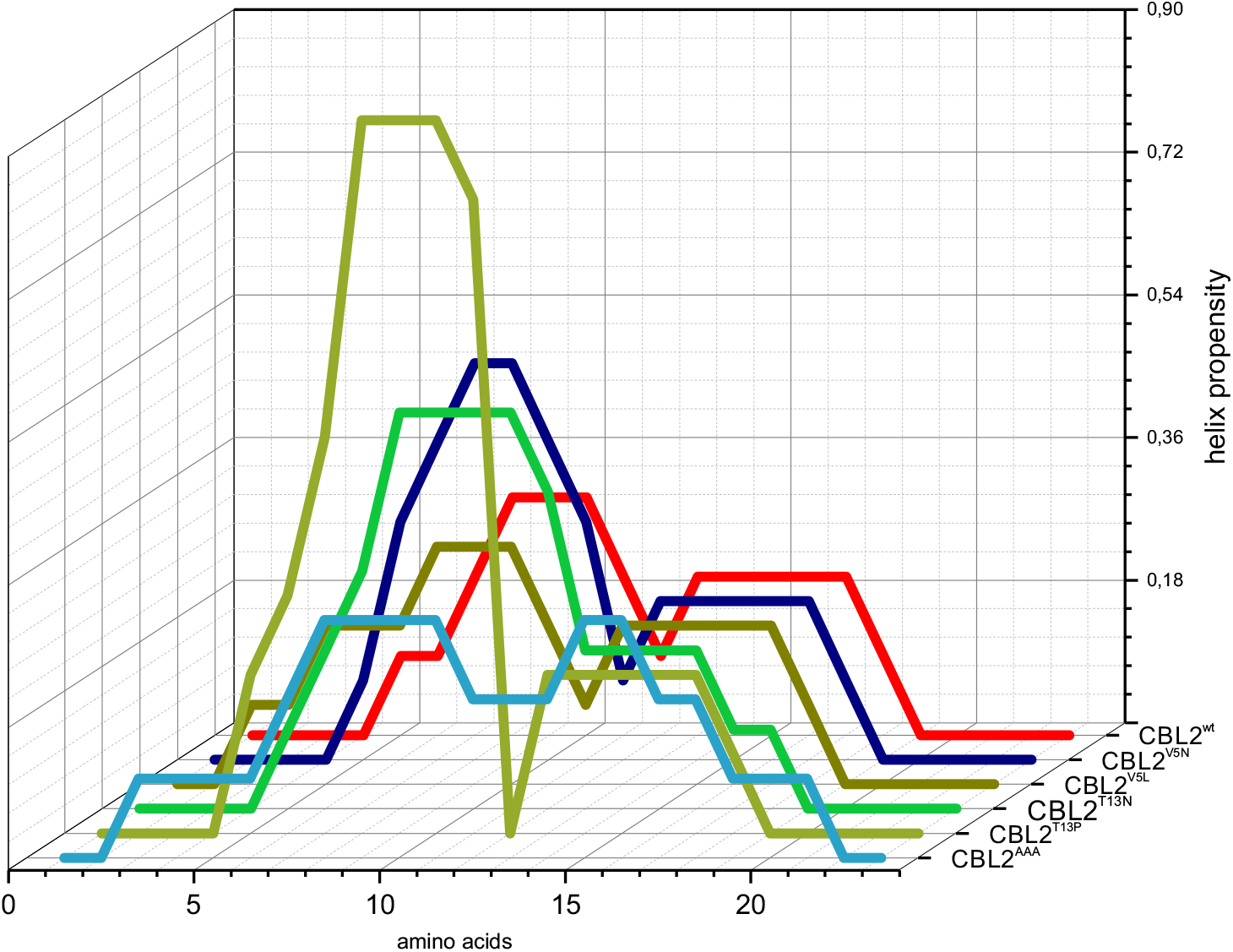
Helix propensity of CBL2 variants used. Helix propensity on the amino acid level was determined using Agadir. Shown are the propensity levels for CBL2^wt^, CBL2^V5N^, CBL2^V5L^, CBL2^T13N^, CBL2^T13P^, CBL2^AAA^.

